# Spatiotemporal variation in honeybee (*Apis mellifera* L.) virome composition across landscape types and seasons

**DOI:** 10.64898/2026.05.27.728278

**Authors:** Rameshwor Pudasaini, Danielle Kroh, Hongmei Li-Byarlay

## Abstract

Honeybees face increasing threats from biotic stress of viral pathogens that can severely impact colony health and contribute to global colony decline. However, comprehensive studies of biotic stress and composition of bee viruses across different environmental contexts and seasons remain scarce. This study aims to characterize and compare the diversity, abundance and composition of the *Apis mellifera* L. virome across three different landscapes (conventional, organic, and roadside) and seasonal gradients (early vs. late season) to better understand how environmental and temporal factors affect viral communities in honeybees. *A. mellifera* were collected from three different habitats (conventional farm, organic farm, and roadside habitat) during the spring and summer of 2024. Total RNA was extracted individually from whole honeybees and mRNA libraries were prepared, which were subsequently used for sequencing on an Illumina NovaSeq X Plus platform using paired-end 150 bp reads. Several bacteriophages, putative novel viruses, plant-, insect- and bee-associated viruses were detected in the honeybee viromes including Sacbrood virus, Black queen cell virus, Deformed wing virus-B (previously known as Varroa destructor virus-1) and Deformed wing virus. Furthermore, both habitat types and seasons influence viral abundance as majority of detected viruses showed higher abundance in the conventional farm and late season samples. The present findings provide novel insights into the ecological and seasonal dynamics of honeybee-virus interactions and contribute to strategies for improving honeybee health and resilience.

## Introduction

Honeybees are considered as the most important manage pollinators due to their large colony size, transportability, foraging efficiency, and visiting a single crops species during a foraging bout^1,2^. As they are responsible for pollinating majority of agricultural crops and their pollination can increase crops yield significantly^6,7^. Their pollination plays a critical role in natural as well as agricultural ecosystems and contributes significantly to global food security^3-5^. In the United States, honeybee is responsible for pollinating more than 130 types of fruits, nuts and vegetables that worth about USD 15 billion annually^8^.

Globally, honeybee populations have experienced substantial declines over the past decades^9^. Both biotic and abiotic stressors, including habitat loss, pesticide exposure, nutritional stress, parasites, and pathogens, contribute to this decline^10,11^. Among the biotic stressors, viruses are considered as one of the most widespread and significant threats to honeybee health^12,13^. Viral infection can negatively affect individual bee physiology, behavior, foraging efficiency and survival, ultimately contributing to colony collapse^14,15^. In addition, virus infected honeybee colonies may experience reduce in workers lifespan, brood mortality and queen failure^12,16^. They also impair immune system of honeybees and cause developmental deformities such as wing deformities due to Deformed wing viruses (DWV)^17^. In addition, viruses often interact with other stressors, such as poor nutrition, pesticides exposure or ectoparasite (*Varroa destructor*) creating synergistic effects that further weaken honeybee colonies^18-20^.

More than 70 viruses have been identified in honeybees, most of which are positive-sense single-stranded RNA viruses^21^. Among them, DWV-A, DWV-B, Acute bee paralysis virus (ABPV), Israeli acute paralysis virus (IAPV), Black queen cell virus (BQCV), Chronic bee paralysis virus (CBPV) and Sacbrood Virus (SBV) are considered the most pathogenic and widely distributed^12,22^. These common bee viruses belong to different families with DWV and SBV belong to family Iflaviridae, BQCV, ABPV, and IAPV belong to family Dicistroviridae; however, CBPV remains unclassified^23-26^. However, a recent study reported several novel DNA bee-associated viruses belongs to Parvoviridae family^27^. The transmission and replication of many honeybee viruses are further facilitated by parasitic mites such as *V. destructor*, which act as efficient vectors and amplify viral infections within colonies^12,17^.

Environmental heterogeneities due to variation in habitat types and temporal variation across seasons may be the fundamental ecological drivers of honeybee virus composition and other pathogens. Worker honeybees forage across diverse environments with varying floral resources, species interactions, and management practices^28,29^. These environmental heterogeneities can structure viral diversity and community composition in honeybee colonies by modulating nutrient availability, environmental stressors, interactions with other insect species and pathogen transmission pathways ^30-32^. In addition to spatial variation, seasonal changes may also contribute to shaping viral dynamics by affecting colony demographics, modify interspecific interactions, resource availability, and transmission opportunities^33,34^.

In this study, we characterized the virome of honeybees collected from different landscape types and seasons using RNA sequencing-based metagenomic analysis. Honeybee samples were collected from conventional farm, organic farm and roadside habitat in May and July in the same year. Conventional farms often involve the direct inputs of different agrochemical such as pesticides and fertilizers, intensive agricultural practices and generally monoculture, which may reduce floral diversity and resource availability for pollinators^35,36^. In contrast, organic farms represent no synthetic agrochemicals and provide higher plant diversities that reduce pollinator overlaps and limit viral persistence in the environment^35^. Roadside habitats which are generally unmanaged systems and typically have minimal exposure to agrochemicals, but they may experience higher levels of traffic-related pollutants^37,38^. Seasonal variation was defined by two sampling periods representing May as an early season characterized by spring conditions with higher precipitation variability and increasing floral resources and July as a late season characterized by mid-summer conditions with higher temperatures, relatively more stable weather with occasional convective rainfall. Understanding how these contrasting habitats and seasonal variation shape the honeybee viromes provide critical insights into pathogen dynamics and foundation to improve colony health and resilience. The present study aimed to identify viruses associated with honeybees (*A. mellifera*) and compare their abundance across different seasons and landscape management types using metagenomic studies. Based on this, we hypothesized that both seasons and landscape management types would influence the diversity and composition of honeybee-associated viruses.

### Materials and Methods Sample collection

Two honeybees (*A. mellifera*) hives with identical numbers of workers, brood, honey and pollen stores were installed at each collection site (WB as conventional farm, Woo as organic farm and Gas as roadside habitat) and forager honeybees were collected using sweep nets in May and July of 2025 (Supplementary Table S1. More than 50 individual honeybees were collected from each collection site. Collected honeybees were immediately flash-frozen in liquid nitrogen and stored at −80 °C until used for RNA extraction. For each collection site and season, 4 honeybee samples were individually used for RNA extraction. A total of 24 honeybee samples were used for sequencing from three collection sites. The details about the collection sites, seasonal and landscape distribution of honeybee samples are presented in Supplementary Table S1.

### RNA extraction, library preparation and Illumina sequencing

Total RNA was extracted using whole honeybee samples using the RNeasy Mini Kits (Qiagen, Hilden, Germany) with following the manufacturer’s protocol. Genomic DNA (gDNA) eliminator spin columns (Qiagen, Hilden, Germany) were applied to remove genomic and environmental DNA. Before further procced for mRNA library preparation, RNA concentration and purity were measured using a NanoDrop spectrophotometer (Thermo Fisher Scientific, USA) and RNA integrity was determined using an Agilent 4200 TapeStation system (Agilent Technologies, USA).

mRNA libraries were prepared using Qiagen’s QIAseq Stranded mRNA Enrichment Kit (Qiagen, Hilden, Germany) in combination with the QIAseq Low Input RNA Library Kit (Qiagen, Hilden, Germany), employing both N6-T RT and ODT-RT primers for reverse transcription. A total of 200□ng of input RNA per sample, quantified using a Qubit fluorometer (Thermo Fisher Scientific, USA), was used for library construction. RNA fragmentation was performed for 3□minutes, followed by first- and second-strand synthesis and adapter ligation according to the manufacturer’s protocol. Unique dual indexes were incorporated using the QIAseq UX96 Index Kit UDI-B (Qiagen, Hilden, Germany). Libraries were amplified for 18 PCR cycles, purified, and assessed for quality prior to sequencing using both Qubit Fluorometer (Thermo Fisher Scientific, USA) and Agilent 4200 TapeStation system (Agilent Technologies, USA). Final libraries were pooled and sequenced on an Illumina NovaSeq X Plus platform (Illumina Ibc., San Diego, CA, USA) using paired-end 150 bp reads to generate high-depth mRNA expression data.

### Data cleaning, virome assembly and viral identification

The raw reads underwent quality control and trimming using Trimmomatic (v0.39) to remove low-quality bases and adapter sequences^39^. Further, the host derived reads were removed using Bowtie2 (v2.5.1) by aligning to the honeybee reference genome Amel_HAv3.1 (GCF_003254395.2, NCBI RefSeq)^40^ and the resulting Sam files were processed using SAMtools (v1.17) for conversion to BAM format, sorting, and indexing^41^. The remaining reads were then assembled using de novo assembly with MEGAHIT (v1.2.9) to reconstruct longer contigs^42^.

BLAST-PLUS (v2.16.0) was used to identify virus sequences to search for assembled contigs to default nr/nt database with E-value cutoff of 1e-7^43^ with the blast results of contigs >500 nt. A contig was assigned as a known virus if it aligned stringently (E = 0.0 and percent identity: >90%) to existing viruses. If the alignment is less stringent (0 < E-value < 1e-7 and percent identity: 69 □ <90%), it was assigned as a putative novel virus.

### Relative abundance of viruses and differential analysis between landscapes

Raw mapped read counts were normalized as counts per million mapped reads (CPM) to account for differences in sequencing depth among libraries. CPM values were calculated as

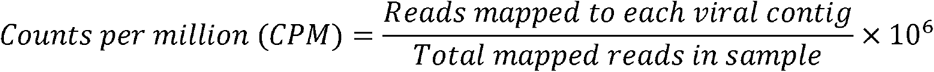

Normalized CMP values were used as a relative measure of viral abundance across samples and samples were grouped based on landscape types and seasons.

## Results

### Reconstruct virome from viral RNA-Seq reads

Using MEGAHIT, we assembled more than 21,857 contigs at least 500 base pairs. With nucleotide BLAST, we obtained 639 BLAST hits, which resulted in the identification of 296 viral contigs. Among these, 95 contigs corresponded to eukaryotic viruses after excluding bacteriophages. We further classified these viral sequencing based on similarity to known viruses. 59 contigs showed exact matches (E = 0.0 and percent identity: >90%) with known viruses, and remaining 237 contigs were considered as either putative novel viruses (0 < E-value < 1e-7 and percent identity: 69 □ <90%) or other virus related contigs. Overall, our metatranscriptomic analysis identified a diverse set of viral sequences, including both known viruses, and several putative and viral related contigs.

### Relative abundance of bacteriophage, giant virus, and environmental viruses

The abundance of bacteriophage, giant virus and environmental viruses detected in honeybee samples across collection sites and seasons is presented in Figure 1. The heatmap was created based on log_2_-transformed CPM values. A total of 16 viruses were detected with 4 bacteriophages, 5 giant viruses, and reaming environmental, protist or animal viruses with differences in abundance across collection habitats and seasons. However, some viruses, such as Staphylococcus phage Andhra, detected with high abundance across all collection habitats and seasons. Some viruses exhibited site-specific distributions; for example, Escherichia phage 500465-1 was detected only in organic farm in very low abundance.

**Figure 1.**
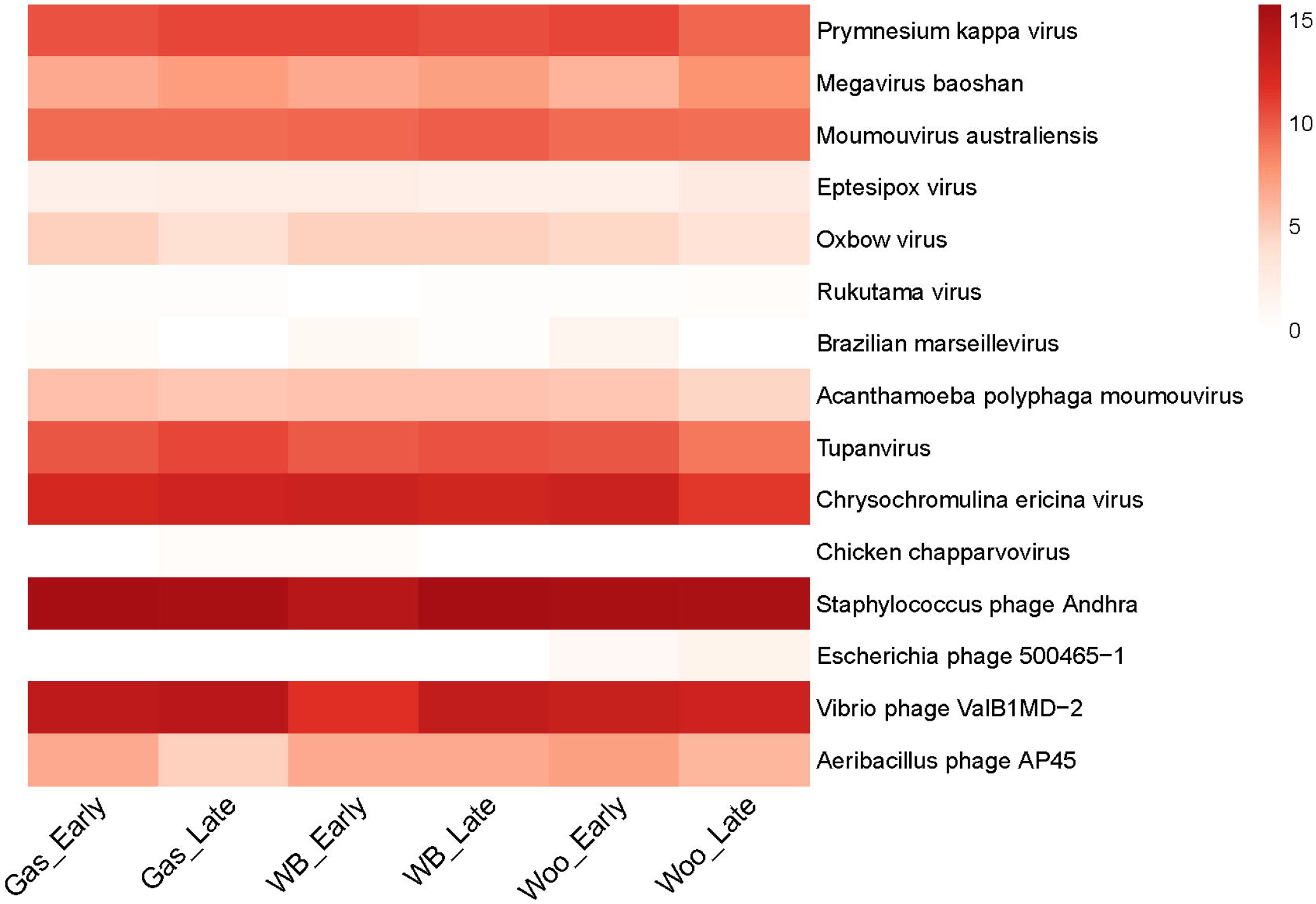
Heatmap of log_2_ transformed counts per million (CPM) values of bacteriophage and environmental viral abundance detected in honeybee samples across locations and seasons. Gas-Early: Roadside habitat in May; Gas-Late: Roadside habitat in July; WB-Early: Conventional farm in May; WB-Late: Conventional farm in July; Woo-Early: Organic farm in May; Woo-Late: Organic farm in July.

### Putative novel viruses

Identified putative novel viruses detected in *A. mellifera* viromes are presented in Table 1. These viral contigs showed significant similarity to known viruses; however, were not exact matches with them (0 < E-value < 1e-7 and percent identity: 69 □ <90%), suggesting the presence of putative novel viruses. A total of 15 putative novel viruses were detected with the closest viral homolog to insects, plants, protists and algae. For example, closest to an insect virus, Choristoneura biennis entomopox virus detected with complete genome, 76.29% of identity and 7.71E-25 of E value. Similarly, closet to a plant virus, Tomato ringspot virus detected with complete sequence, 84.71% of identity and 2.82E-176 of E value.

**Table 1.** Identified putative novel viruses detected in *A. mellifera* virome assemblies assembled from RNA-Seq data with 0 < E-value < 1e-7 and percent identity 69-89%.

| Contig ID | Closest viral homolog | Assembly Level | Percent identity | Alignment length (bp) | E-value | Potential host |
| --- | --- | --- | --- | --- | --- | --- |
| k141_11377 | Moumouvirus australiensis | Complete genome | 81.82 | 121 | 4.19E-19 | Protists |
| k141_16188 | Chrysochromulina ericina virus | Complete genome | 82.10 | 229 | 1.12E-47 | Algae |
| k141_16785 | Tupanvirus | Complete genome | 74.53 | 373 | 6.08E-32 | Protists |
| k141_16925 | Choristoneura biennis entomopoxvirus | Complete genome | 76.29 | 232 | 7.71E-26 | Insect |
| k141_17951 | Tomato ringspot virus | Complete sequence | 84.71 | 628 | 2.82E-176 | Plant |
| k141_18315 | Acanthamoeba polyphaga moumouvirus | Complete genome | 80.92 | 173 | 1.57E-29 | Protists |
| k141_18410 | Brazilian marseillevirus | Complete genome | 80.19 | 212 | 3.66E-37 | Protists |
| k141_1864 | Prymnesium kappa virus | Chromosome: 1 | 81.66 | 229 | 2.03E-45 | Algae |
| k141_19092 | Actinidia virus X | Complete cds | 84.70 | 366 | 1.10E-99 | Plant |
| k141_20496 | Tupanvirus soda lake | Partial genome | 72.01 | 718 | 5.48E-44 | Protists |
| k141_20549 | Grapevine Bulgarian latent virus | Complete genome | 80.12 | 327 | 5.34E-58 | Plant |
| k141_20734 | Cafeteria roenbergensis virus | Complete genome | 78.28 | 221 | 7.84E-31 | Protists |
| k141_25936 | Soybean latent spherical virus | Complete sequence | 78.53 | 624 | 7.56E-104 | Plant |
| k141_4490 | Powai lake megavirus | Complete genome | 77.57 | 214 | 5.26E-26 | Protists |
| k141_7299 | Moumouvirus australiensis | Complete genome | 69.64 | 1535 | 1.36E-45 | Protists |

### Plant-associated viruses

A total of 19 plant viruses were detected in *A. mellifera* viromes samples collected from different habitats and seasons (Table 2). The identified viruses represent diverse taxonomic groups and include well-characterized species including Soybean carla virus 1 (SCV1), White clover cryptic virus (WCCV), Red clover cryptic virus (RCCV) and Cherry leaf roll virus (CLRV).

**Table 2.**
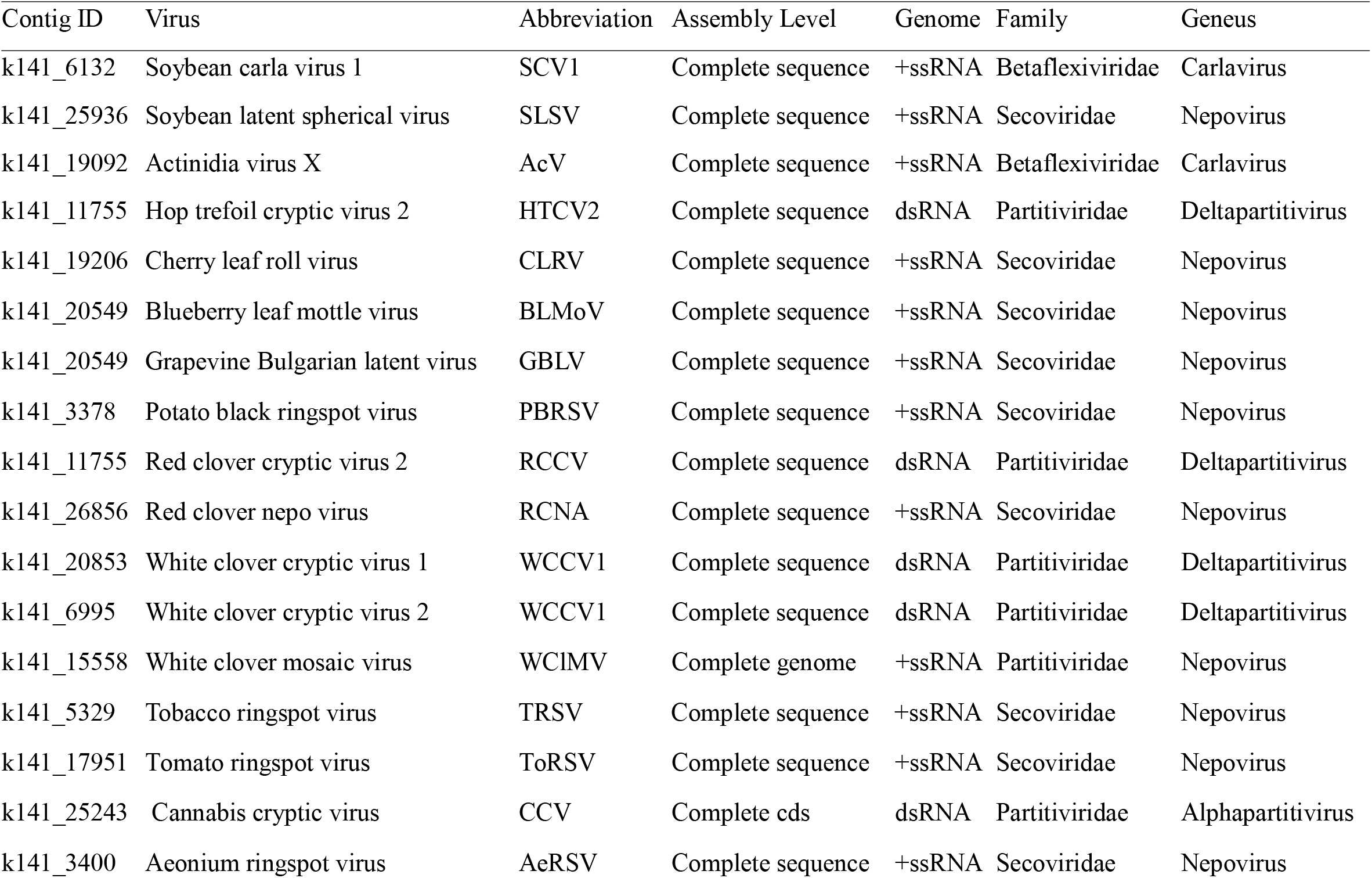
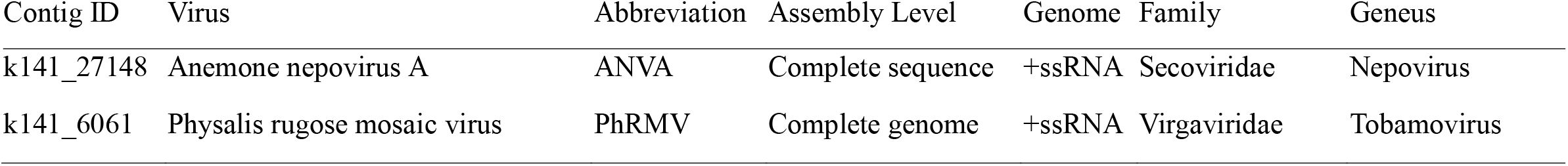
Identified plant viruses detected in *A. mellifera* virome assemblies assembled from RNA-Seq data with E = 0.0 and 0percent identity more than 90%.

### Bee and other insect associated viruses

Identified bee and insect viruses detected in *A. mellifera* viromes collected from the conventional farm, organic farm and roadside habitat are presented in Table 3. A total of 11 bee and other insect-associated viruses were identified across the analyzed honeybee samples, including 4 bee-associated viruses and 7 other insect-associated viruses. Some of the identified insect-associated viruses were Choristoneura fumiferana granulovirus, Diolcogaster facetosa bracovirus, Drosophila innubila nudi virus, Blattodea nairo-related virus, Aphis citricidus bunyavirus and Shamonda Virus. Furthermore, SBV, BQCV, DWV and DWV-B (previously known as Varroa destructor virus-1) bee-associated viruses were detected.

**Table 3.** Identified bee and insect viruses detected in *A. mellifera* virome assemblies assembled from RNA-Seq data with E = 0.0 and percent identity more than 90%.

| Contig ID | Virus | Abbreviation | Assembly Level | Genome | Family | Geneus | Host |
| --- | --- | --- | --- | --- | --- | --- | --- |
| k141_14135 | Sacbrood virus | SBV | Complete genome | +ssRNA | Iflaviridae | Iflavirus | Bee |
| k141_6297 | Black queen cell virus | BQCV | Complete cds | +ssRNA | Dicistroviridae | Triatovirus | Bee |
| k141_21979 | Black queen cell virus-B | BQCV-B | Complete genome | +ssRNA | Iflaviridae | Iflavirus | Bee |
| k141_9247 | Deformed wing virus | DWV | Complete genome | +ssRNA | Iflaviridae | Iflavirus | Bee |
| k141_15254 | Choristoneura fumiferana granulovirus | CfGV | Complete sequence | dsDNA | Baculoviridae | Betabaculovirus | Insect |
| k141_4403 | Diolcogaster facetosa bracovirus | DfBV | Complete sequence | dsDNA | Polydnaviridae | Bracovirus | Insect |
| k141_23911 | Drosophila innubila nudi virus | DiNV | Complete sequence | -ssRNA | Nudiviridae | Alphanudivirus | Insect |
| K141_4698 | Blattodea nairo-related virus | OKIAV321 | Complete sequence | -ssRNA | Nairoviridae | Othonairovirus | Insect |
| k141_26160 | Aphis citricidus bunyavirus | AcBV | Complete sequence | -ssRNA | Phenuiviridae | Orthobunyavirus | Insect |
| k141_14135 | Hubei picorna-like virus 40 | HPLV40 | Complete cds | +ssRNA | Unclassified | Unknown | Insect |
| k141_4211 | Shamonda Virus | SHAV | Complete sequence | -ssRNA | Peribunyaviridae | Orthobunyavirus | Insect |

### Relative abundance of viruses and differential analysis between landscapes

The abundance of plant and insect viruses detected in honeybee samples across three collection sites and two seasons are presented in Figure 2 a,b. The heatmap was created based on log_2_-trasformed CPM values. Several viruses showed variation in their abundance across locations and seasons. Comparing the collection sites, the higher viral abundance was observed in the honeybee samples collected from conventional farms compared to organic farms and roadside habitat samples for most of the viruses. In addition, early-season samples also showed higher viral abundance than late season samples. For example, BQCV and DWV-B were presented in higher abundance in conventional farm and roadside habitat than organic farm in late season samples. However, SBV was presented at a relatively higher level in late season samples of organic farm.

**Figure 2.**
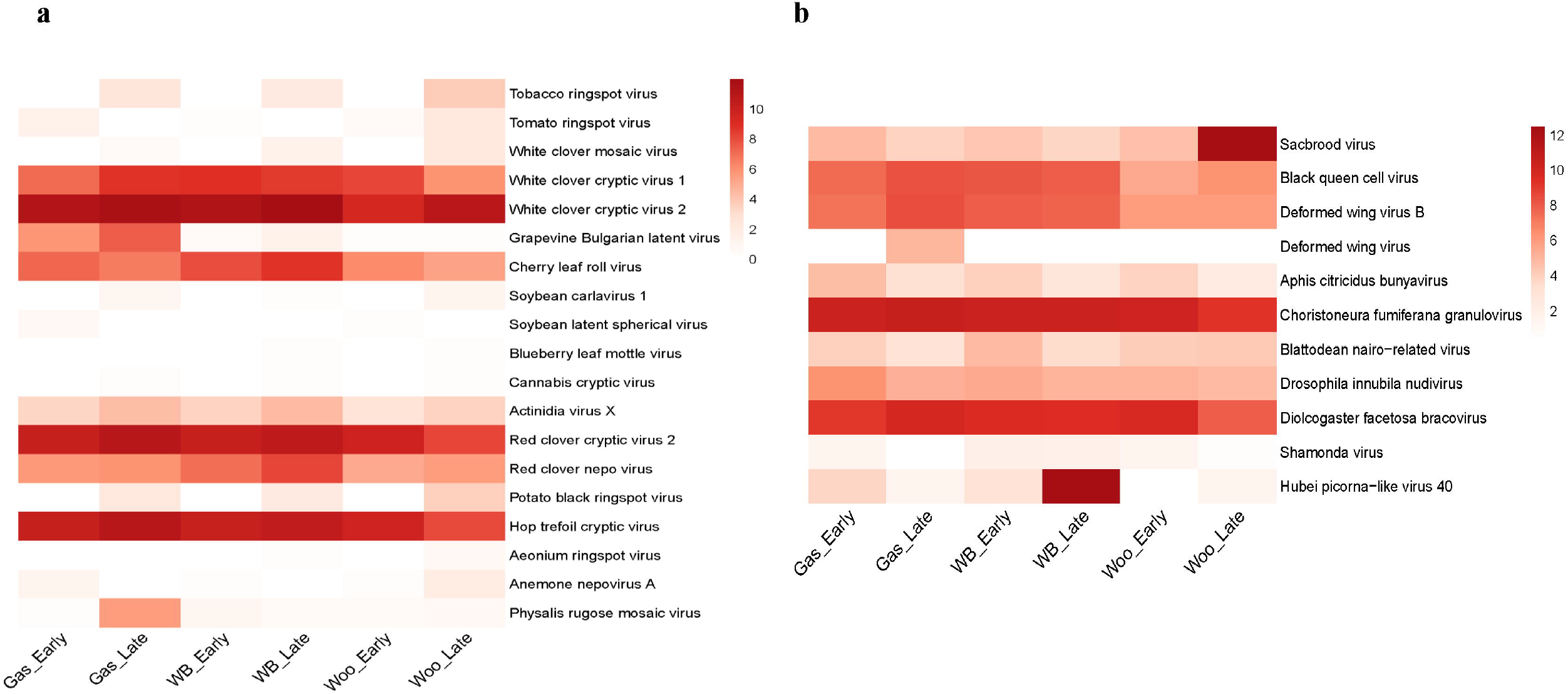
Heatmap of log_2_ transformed counts per million (CPM) values of (a) plant-associated and (b) insect-associated viral abundance detected in honeybee samples across locations and seasons. Gas-Early: Roadside habitat in May; Gas-Late: Roadside habitat in July; WB-Early: Conventional farm in May; WB-Late: Conventional farm in July; Woo-Early: Organic farm in May; Woo-Late: Organic farm in July.

## Discussion

This study provides compositions of the honeybee virome across different landscape types and seasonal conditions. In addition to known bee-associated viruses, bacteriophages, environmental viruses, putative novel viruses, plant- and insect-associated viruses were detected in the present study, which show the broad exposure of honeybees to diverse viral communities during foraging^47^. Furthermore, based on the present findings, environmental conditions influence honeybee viral abundance as higher viral abundance was observed in the conventional farm and late season collected honeybee samples.

The detection of bacteriophages and environmental viruses suggests that honeybees are exposed to viruses not only through direct infection but also through environmental contact while foraging, social interaction and their hives^47,48^. Other studies also detected different bacteriophages in honeybee virome^47,48^. The bacteriophages in honeybee viromes may originate from honeybee-associated gut microbial communities^49^.These bacteriophages viruses are generally non-pathogenic to honeybees and play a crucial role in maintaining gut health, regulating bacterial populations and influencing colony immunity^47^. Indeed, bacteriophages can be used as a potential biological agent to treat different bacterial diseases, such as foulbrood disease, *Paenibacillus larvae* in honeybee^50,51^.

Detection of putative novel viruses show the complexity and underexplored of the honeybee virome compositions. These findings indicate that the honeybee-associated virome may harbor many unexplored viral communities^52^. Other different studies also reported novel and unknown viruses in honeybees viromes^27,53,54^. However, further genomic and phylogenetic studies are necessary to confirm their novelty, host specificity, biological relevance and potential impacts on honeybee health^52^. Expanding viral reference genomes in honeybees virome associated with environmental sources will be important for improving future taxonomic resolution.

Several plant-associated viruses were detected in honeybee viromes in the present study. Other studies also frequently detected plant-associated viruses in honeybee viromes^55,56^. A study also reported 15 plant-associated viruses in honeybee viromes^55^. These plant-associated viruses in the honeybee virome may be detected into honeybee samples through pollen and nectar during foraging, because many plant viruses can travel with attaching pollen grains^57^. Overall, the detection of plant viruses in the honeybee viromes indicates that honeybees may act as carrier of plant viruses in agricultural ecosystems^58^.

Similarly, a total of 7 insect-associated viruses were detected in the present study. The detection of these insect-associated viruses suggests potential viral spillovers due to complex interactions within pollinators, pests, parasitoids and other flower visiting arthropods within pollinators’ ecosystems^59^. Understanding these connections could clarify how ecological communities influence viral persistence and spread. This may also reflect shared viral communities and transmission pathways through shared floral resources, potential cross-species virus transmission or passive carriage of viral particles by honeybees^60^.

We detected 4 important bee-associated viruses in honeybee viromes collected from different landscape types and seasons. All detected 4 bee-associated viruses in this study that include SBV, BQCV, DWV and DWV-B (previously known as Varroa destructor virus-1) are commonly reported in honeybee viromes and associated with diseases in honeybees. A study also reported 9 bee-associated viruses in honeybee viromes^55^. The presence of bee-associated viruses indicates the host-specific viruses that may directly be associated with honeybee health, and may also represent both latent and active infections within colonies^13^. However, their presence does not necessarily mean active infections to honeybees^61^, but interactions with other stressors such as nutritional limitation, pesticide toxicity and parasitic infections may become more pathogenic and contribute to colony health. In addition, there will be always risk of transmission of these bees associated with viruses through social interaction such as trophallaxis, grooming, and brood caring within colonies or other colonies through vectors like *Varroa* mites.

Variation in management practices, floral diversity, and environmental stressors in different landscape types play a critical role in shaping honeybee viral compositions^20^. The present results suggest that habitat types influence viral abundance in honeybees as higher viral abundance was observed in the conventional farm and lower in the organic farm. In conventional farms, certain agricultural practices such as agricultural intensification and monoculture often reduce floral diversity and resource availability for pollinators, confining them to limited floral resources^35,36^. Consequently, it facilitates horizontal viruses’ transmission within pollinators and influences viral composition and infection^62^. Diverse landscapes, such as in organic farms, may dilute viral transmission by supporting greater ecological diversity, heterogeneous foraging environments, reducing pollinator overlaps, and limiting viral persistence in the environment^35^. Roadside habitats, on the other hand, are often characterized by rich floral resources; however, atherogenic disturbances such as traffic-related pollution and habitat fragmentation, which may also influence viral diversity and transmission dynamics^37,38^.

Seasonal variation also contributes to honeybee virome compositions with differences in environmental conditions, floral resource availability, and other insect activities^20,63^. In the present study, higher viral abundance was observed in late season honeybee samples. Early seasonal conditions, such as May, are characterized by emerging floral resources and lower foraging activities, whereas mid-summer conditions, such as July, are associated with higher vegetation, higher foraging activities, and greater interspecific or intraspecific interactions. Such seasonal dynamics may enhance opportunities for virus transmission among pollinators and between plants and insects, thereby influencing viral abundance and diversity in honeybees^33^.

Unexpectedly high viral abundances were detected in some collection sites for some viruses, for instance, SBV showed exceptionally high abundance in late-season samples from the organic farm. These inconsistences may be due to insufficient sample size. Therefore, future studies should include higher number of sampling sites and increased sample sizes to reduce the influence of outliers in viral distribution.

The presence of diverse viral groups reflects extensive environmental exposure in honeybee virome; however, it does not necessarily mean to active infection or pathogenicity. The functional and biological significance of these detected viruses remains unclear. Therefore, future studies on host immune responses and influences on honeybee health are necessary for proper understanding of their ecological and biological relevance. Overall, this study demonstrates that both landscape and seasonal variation shape honeybee virome compositions and abundance. The present study findings improve understanding of honeybee viromes composition associated with their ecosystem and implications for improving honeybee health.

## Acknowledgement

We acknowledge the Genomics Shared Resource (GSR) at OSUCCC-The James for their support with the library preparation and sequencing. GSR is funded in part by the National Cancer Institute P30 CA016058. We thank Heaven Strachan for her help in collecting bee samples in the field.

## Funding

This research was supported by USDA NIFA Capacity Building Grant 2021-38821-34576, and USDA NIFA Evans-Allen Fund NI251445XXXXG001.

## Authorship contributions

HL-B-Conceptualization, Methodology, Funding acquisition, Visualization, Project administration, Supervision, Writing - review & editing. DK - Methodology. RP- Methodology, Investigation, Data analysis, Writing - original draft.

## Data availability

The raw reads of RNA-Seq of the present study are available in NCBI BioProject PRJNA1456520.

## Conflict of interest

The authors declare no conflict of interest.

## Supplementary materials

Supplementary material associated with this study are available in the online version of this paper.

## Notes

### Competing Interest Statement

The authors have declared no competing interest.

